# RNA-seq and differential expression analysis of the duck transcriptome: The effect of short-term cage-rearing

**DOI:** 10.1101/2021.05.13.444049

**Authors:** Biao Chen, Wenjie Fang, Yankai Li, Ting Xiong, Mingfang Zhou, Lei Wan, Qiuhong Liu, Wenyan Zhang, Xiaolong Hu, Huirong Mao, Sanfeng Liu

## Abstract

Ducks are an important source of meat and egg products for human beings. In China, duck breeding has gradually changed from the traditional floor-water combination system to multilayer cage breeding. Therefore, the present study collected the hypothalamus and pituitary of 113-day-old ducks after being caged for 3 days, in order to investigate the effect of cage-rearing on the birds. In addition, the same tissues (hypothalamus and pituitary) were collected from ducks raised in the floor-water combination system, for comparison. Thereafter, the transcriptomes were sequenced and the expression level of genes were compared. The results of sequencing analysis showed that a total of 506 and 342 genes were differentially expressed in the hypothalamus and pituitary, respectively. Additionally, the differentially expressed genes were mainly enriched in signaling pathways involved in processing environmental information, including ECM-receptor interaction, neuroactive ligand-receptor interaction and focal adhesion. The findings also showed that there was a change in the alternative splicing of genes when ducks were transferred into the cage rearing system. However, there was no difference in the expression of some genes although there was a change in the expression of the isoforms of these genes. The findings herein can therefore help in understanding the mechanisms underlying the effect of caging on waterfowl. The results also highlight the gene regulatory networks involved in animal responses to acute stress.

## Introduction

The consumption of poultry products across the globe has increased over the years. Notably, China is the world’s largest producer and consumer of duck products (http://www.fao.org/faostat/en/#data/QA/visualize). In addition, the floor-water combination system is the conventional method of rearing ducks in China (Figure 1A). This feeding system is highly dependent on the water area. However, the conventional floor-water combination system of rearing ducks has many shortcomings. Such include poor utilization of water resources, serious pollution to the water environment which limits large scale production as well as disease control and low economic benefits. Therefore, it is necessary to adopt an environmentally friendly breeding model for ducks. Over the recent years, the cage rearing system has developed rapidly in duck production because it is environmentally friendly and has several economic benefits(Bai *et al.* 2020). Moreover, the cage rearing system for ducks is characterized by high stocking densities, clean egg surfaces and high uniformity of waterfowl (Figure 1 B). However, the system has not been explored extensively.

**Figure 1.**
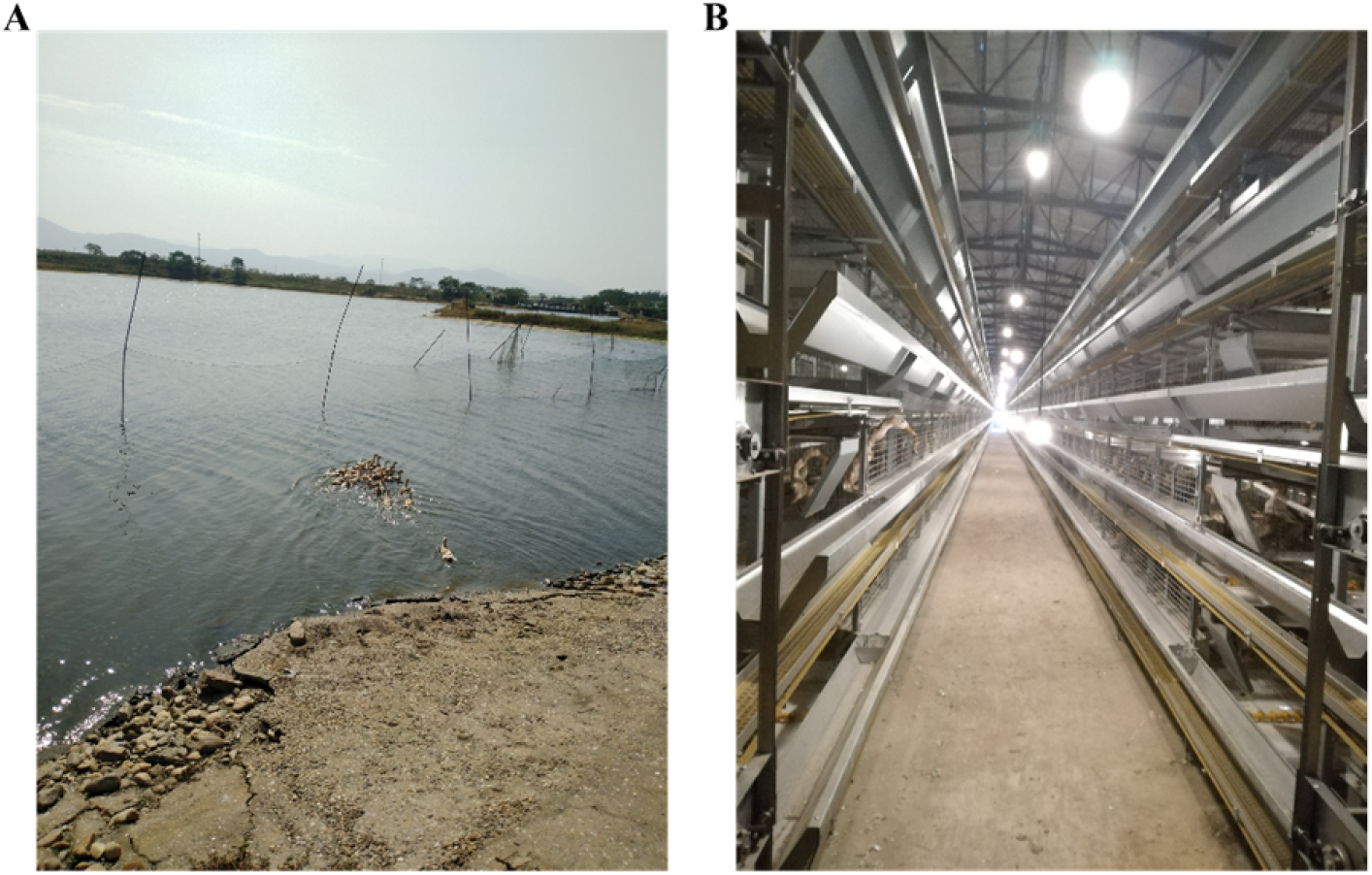
The two feeding systems used in the current study. (**A**) The conventional floor-water combination system for ducks. (**B**) The multilayer cage-rearing system for ducks.

Compared to chicken husbandry, such factors as body size, dabbling, fecal ejection and other habits, complicate the design, manufacture and production of ducks. Additionally, the acute and chronic stress caused by cage rearing are also crucial issues that need to be addressed by duck producers(Zhang *et al.* 2019). In the cage rearing system, the high-density husbandry causes spatial stress to the ducks. Moreover, high density stocking increases the concentration of hydrogen sulfide, ammonia and other harmful gases in duck cages, which are detrimental to the growth and reproduction of the birds(Shepherd *et al.* 2015; Zhao *et al.* 2016). In addition, the air velocity and relative humidity in the cage rearing system are quite different from those in the floor-water combination system(En-cai *et al.* 2020). These environmental changes therefore trigger stress responses and adaptive changes in ducks.

Moreover, ducks experience several changes in their living environment when they are initially transferred into cages, resulting to stress responses in their bodies(Zhang *et al.* 2019). Stress refers to a non-specific response of the body to external stimuli and can be classified as either acute or chronic stress. Notably, the body mainly responds to stress through the Hypothalamus-pituitary-adrenal axis (HPA)(Nicolaides *et al.* 2014; Oyola and Handa 2017). After the center of the brain receives external stimulation, the information is transmitted to the hypothalamus. The hormone secreted from the hypothalamus then stimulates the pituitary gland to secrete the adrenocorticotropic hormone, putting the body in a state of full mobilization. The body in turn experiences an increase in heart rate, blood pressure, body temperature, muscle tension, metabolic level and other significant changes, so as to enhance body activity and cope with emergencies(Munakata 2018; Nicolaides *et al.* 2014).

Therefore, it is important for duck producers and researchers to analyze the internal mechanism of high-density cage-rearing stress in egg laying ducks in order to develop methods for stress alleviation and improve the reproductive performance of cage-reared waterfowl. Consequently, the current study compared the transcriptomes of the hypothalami and pituitary from ducks reared in the two husbandry systems (the traditional floor-water combination and cage-rearing systems). In addition, the study investigated the candidate genes and signaling pathways involved in cage-rearing stress. By exploring the regulatory mechanism of cage-rearing stress in egg laying ducks, this study provides a fundamental basis for establishing new methods of solving caging related stress in waterfowl. This in turn promotes the development and popularization of the waterfowl caging technology and also provides more insights on the molecular mechanism of the interaction between genes and the environment.

## Materials and Methods

### Ethical statement

All the animal experiments in this study were conducted according to the ethical standards of the Jiangxi Agricultural University (JXAULL-2017002). The ducks were sacrificed painlessly.

### Animals and management

A total of 30 female Shan Ma ducks (*Anas platyrhynchos*) were purchased from the Jiangxi Tianyun duck breeding farm (Nanchang, Jiangxi) and reared in the Anyi duck farm of Jiangxi. All the ducks were then transported to the Anyi duck farm after reaching the age of 100 days, then raised in the floor-water combination system (Figure 1A). At the age of 110 days, fifteen ducks were randomly selected and transferred into three cages (100 × 45 × 50 cm, Figure 1B) while the others were retained in the floor-water feeding system. All the ducks were kept at room temperature and fed with the commercial diet produced by the Nanchang Huada Group (Nanchang, Jiangxi, China). Additionally, the ducks raised in the floor-water system (FW group) received natural illumination while the 15 ducks raised in cages (C group) were subjected to a standard regimen involving 16 h of light. All the ducks remained in good health in the entire duration of the experiment.

### Sample collection and RNA extraction

After 3 days of changing the rearing system, all ducks in the FW and C groups were sacrificed. The hypothalami and pituitaries were then sampled using scissors and tweezers, snap-frozen with liquid nitrogen then kept in −80 °C. Thereafter, total RNA was extracted using the TRIzol reagent (Invitrogen, CA, USA), according to the manufacturer’s instructions. In addition, the purity and quantity of RNA were evaluated using NanoDrop ND-1000 (NanoDrop, DE, USA). Moreover, RNA integrity was assessed through electrophoresis in denaturing agarose gel and Bioanalyzer 2100 (Agilent, CA, USA) with a RIN number >7.0.

### RNA library construction and sequencing

After confirming the quality of RNA, four samples from each group were selected to construct the RNA library. Notably, Poly (A) RNA was obtained from 2 μg of total RNA using the Dynabeads Oligo (dT)25-61005 Kit (Thermo Fisher, CA, USA), with two rounds of purification. Thereafter, the poly (A) product was broken into pieces at 94 °C for 5-7 min using the Magnesium RNA Fragmentation Module (NEB, MA, USA). The RNA fragments were then reverse-transcribed using the SuperScript™ II Reverse Transcriptase (Invitrogen) in order to construct the cDNA library. Moreover, the U-labeled second-stranded DNAs were synthesized using the cDNA library obtained from the previous step, *Escherichia coli* (*E.coli*) DNA polymerase I (NEB), RNase H (NEB) and a dUTP solution (Thermo Fisher). After adaptor ligation, the product was treated with the heat-labile UDG enzyme. Finally, the products were amplified through PCR in the following steps: 95 °C for 3 min; 8 cycles of 98 °C for 15 sec, 60 °C for 15 sec and 72 °C for 30 sec then a final step of 72°C for 5 min. Thereafter, the PCR products were submitted for 2 × 150 bp paired-end sequencing (PE150) on the Illumina Novaseq™ 6000 platform (LC-Bio Technology CO., Ltd., Hangzhou, China), following the manufacturer’s instructions.

### Bioinformatics analysis of RNA-Seq data

Cutadapt (https://cutadapt.readthedocs.io/en/stable/) was used to remove the reads containing adaptors and low-quality bases in order to ease analysis in the next step. Thereafter, HISAT2 (https://daehwankimlab.github.io/hisat2/, version: 2-2.0.4) was used to map the reads to the duck genome (https://www.ncbi.nlm.nih.gov/genome/2793?genome_assembly_id=426073). The mapped reads in each sample were then assembled using StringTie (http://ccb.jhu.edu/software/stringtie/, version: stringtie-1.3.4d), with default parameters. Afterwards, a comprehensive transcriptome was reconstructed with the mapped reads from all the samples, using the gffcompare software (http://ccb.jhu.edu/software/stringtie/gffcompare.shtml, version: gffcompare-0.9.8). In addition, StringTie and ballgown (http://www.bioconductor.org/packages/release/bioc/html/ballgown.html) were employed to evaluate the expression levels of all the transcripts and genes in each sample by calculating the fragments per kilobase of exon per million mapped fragments (FPKM). Moreover, the differentially expressed mRNAs (DEGs) and differentially expressed transcripts (DETs) were selected based on the following criteria; |log2 fold change| ≥ 1 and p ≤ 0.05 using the DESeq2 package in R (http://www.bioconductor.org/packages/release/bioc/html/DESeq2.html). After obtaining the list of DEGs, enrichment analyses, including Gene Ontology (GO) and Kyoto Encyclopedia of Genes and Genomes (KEGG), were performed using the DAVID 6.7 functional annotation tool (http://david.abcc.ncifcrf.gov/), in order to assess the potential processes associated with acute stress during cage-rearing.

### Complementary DNA (cDNA) synthesis and qPCR

The cDNA libraries were constructed from total RNA using the Monad MonScript™ All-in-One Kit with DNase (Biopro, Shanghai, China) and reverse-transcription was conducted following the manufacturer’s instructions. Thereafter, the cDNA libraries were diluted in nuclease-free water at a ratio of 1:4 for qPCR. The expression levels of candidate genes were then quantified through qPCR using the 2 × T5 Fast qPCR Mix (TsingKe, Beijing, China) on an ABI QuantStudio 5 system (Thermo Fisher, Waltham, MA, USA). The *GAPDH* gene was used as the internal reference and the final volume of the qPCR reaction was 20 μl. The following conditions were used for qPCR: 95 °C for 3 min; 40 cycles of 95 °C for 10 s, Tm for 1 min and collection of fluorescence at 65 – 95 °C. Additionally, each sample was analyzed in triplicate and the relative expression levels were calculated according to the comparative 2^-ΔΔCt^ method. The primer pairs used in this study were designed using Primer Premier Version 5 (Premier Biosoft, CA, USA). Information on the qPCR primers for the candidate genes and *GAPDH* is shown in Table 1.

**Table 1.**
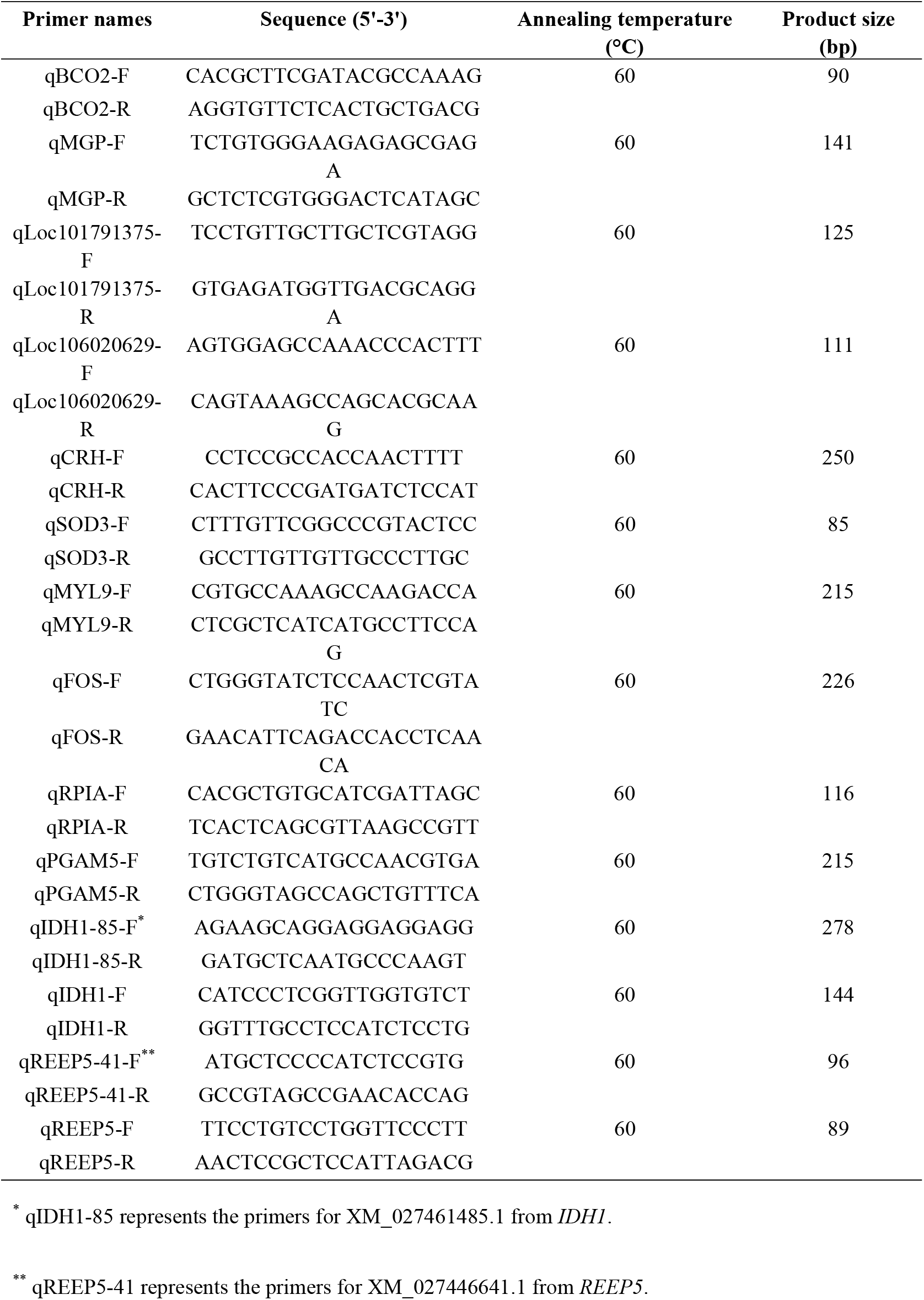
Primer information

### Statistical Analysis

The qPCR results were presented as the mean ± standard error of the mean (S.E.M.). In addition, the Students two-tailed t-test was used for statistical analysis. The level of significance was set at * (p < 0.05), ** (p < 0.01) and *** (p < 0.001).

### Data Availability

Supplemental files available at FigShare. Figure S1: The distribution of mapping region on reference genome, Figure S2: Differential expression analysis of all transcripts, Table S1: Overview of quality of sequencing data, Table S2: All genes detected in current project, Table S3: All the DEGs in the C_H vs. FW_H and C_P vs. FW_P comparisons, Table S4: All information of DETs, Table S5: List of GO enrichment with all DEGs in C_H vs. FW_H, Table S6: List of GO enrichment with all DEGs in C_P vs. FW_P, Table S7: List of KEGG pathways with all DEGs in C_H vs. FW_H, Table S8: List of KEGG pathways with all DEGs in C_P vs. FW_P, Table S9: List of the alternative splicing genes in C_H vs. FW_H and C_P vs. FW_P comparisons. All sequencing data were uploaded to in the Gene Expression Omnibus (GEO) with accession number GSE173134 (https://www.ncbi.nlm.nih.gov/geo/query/acc.cgi?acc=GSE173134).

## Results

### Summary of Transcriptome Data

In this study, a mean of 48,143,316 validated reads were obtained from each sample. In addition, the proportion of validated reads whose base recognition accuracy met the Q30 standard (the error rate was less than 1%) was greater than 97.6% (Table S1).

All the clean reads were then mapped to the duck genome (*Anas platyrhynchos*). The results in Table 2 show that the proportion of mapped validated reads was not less than 74.88% in all the sixteen samples, the percentage of unique mapped reads ranged from 40.44% to 58.50% and the proportion of Pair-end (PE) mapped reads ranged from 53.95% to 78.60%. Moreover, sequence alignment showed that more than 88.3% of the validated reads mapped to the exonic region of each sample (Figure S1). Furthermore, a total of 24,263 genes were detected in all the samples, including annotated and novel genes (Table S2). Out of all the genes, the Growth Hormone 1 (GH1) gene had the highest level of expression after *GAPDH*, in both the hypothalamus and pituitary (Table S2).

**Table 2.**
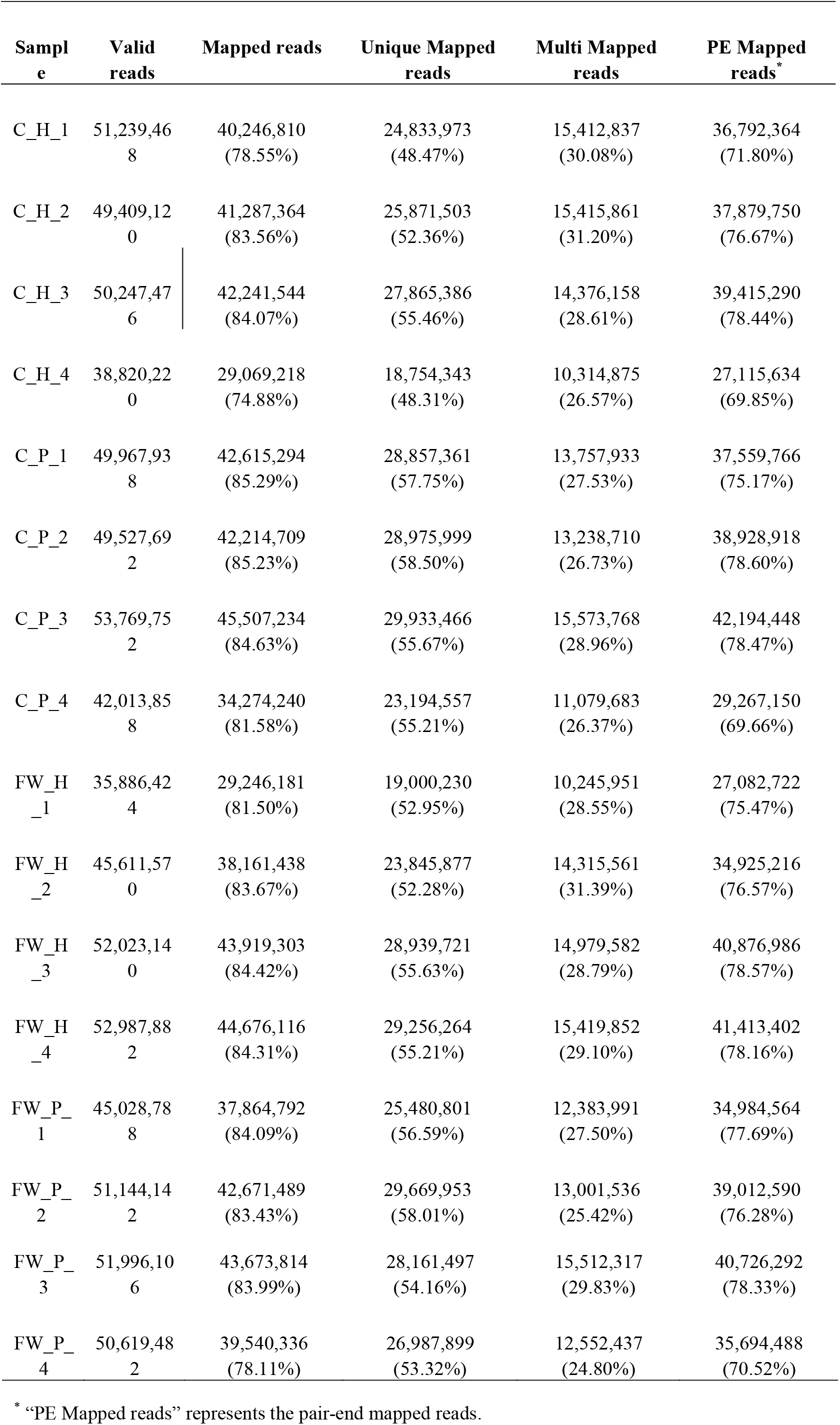
The mapping statistics of validated reads

### Identification of differentially expressed genes (DEGs)

In order to investigate the potential candidate genes and pathways associated with cage-rearing stress, the study performed differential expression analysis, where |log2 fold change| ≥ 1 and p-value ≤ 0.05 were selected as the screening criteria (Figure 2A and B). As a result, a total of 506 DEGs were obtained in the hypothalamus (C_H vs. FW_H) out of which 445 were up-regulated and 61 were down-regulated (Figure 2C). On the other hand, a total of 342 DEGs, including 75 up-regulated and 267 down-regulated genes, were obtained in the pituitary (C_P vs. FW_P, Figure 2C). All the DEGs are listed in Table S3. Notably, the top 3 most highly expressed genes with the smallest q-values in the hypothalamus (C_H vs. FW_H) were *GH1*, Transthyretin (TTR) and Chromogranin A (CHGA). On the other hand, the Nuclear Receptor Subfamily 4 Group A Member 1 (NR4A1), Transgelin (TAGLN) and Superoxide Dismutase 3 (SOD3) were the top 3 most highly expressed genes in the pituitary (C_P vs. FW_P). Additionally, the study assessed the Differentially Expressed Transcripts (DETs) using the same criteria as those for DEGs (Figure S2A and B). A total of 1019 DETs were detected in the hypothalamus (C_H vs. FW_H), out of which 612 were up-regulated and 407 were down-regulated (Figure 2D). Moreover, a total of 916 DETs were obtained in the pituitary (C_P vs. FW_P), out of which 438 were up-regulated and 478 were down-regulated (Figure 2D). All the information on DETs is shown in Table S4. The study then performed clustering analysis on the top 50 DEGs (50 DEGs with the smallest q-values) in each comparison and the findings were presented as the normalized log_2_ FPKM. The results of cluster analysis showed that the upregulated and downregulated genes were quite distinct in each comparison (Figure 2E and F). Clustering analysis was also performed on the top 100 DETs in each comparison. Similar results to those of clustering analysis on DEGs were obtained. However, the number of up- and down-regulated DETs were almost similar, contrary to that of DEGs (Figure S2C and D).

**Figure 2.**
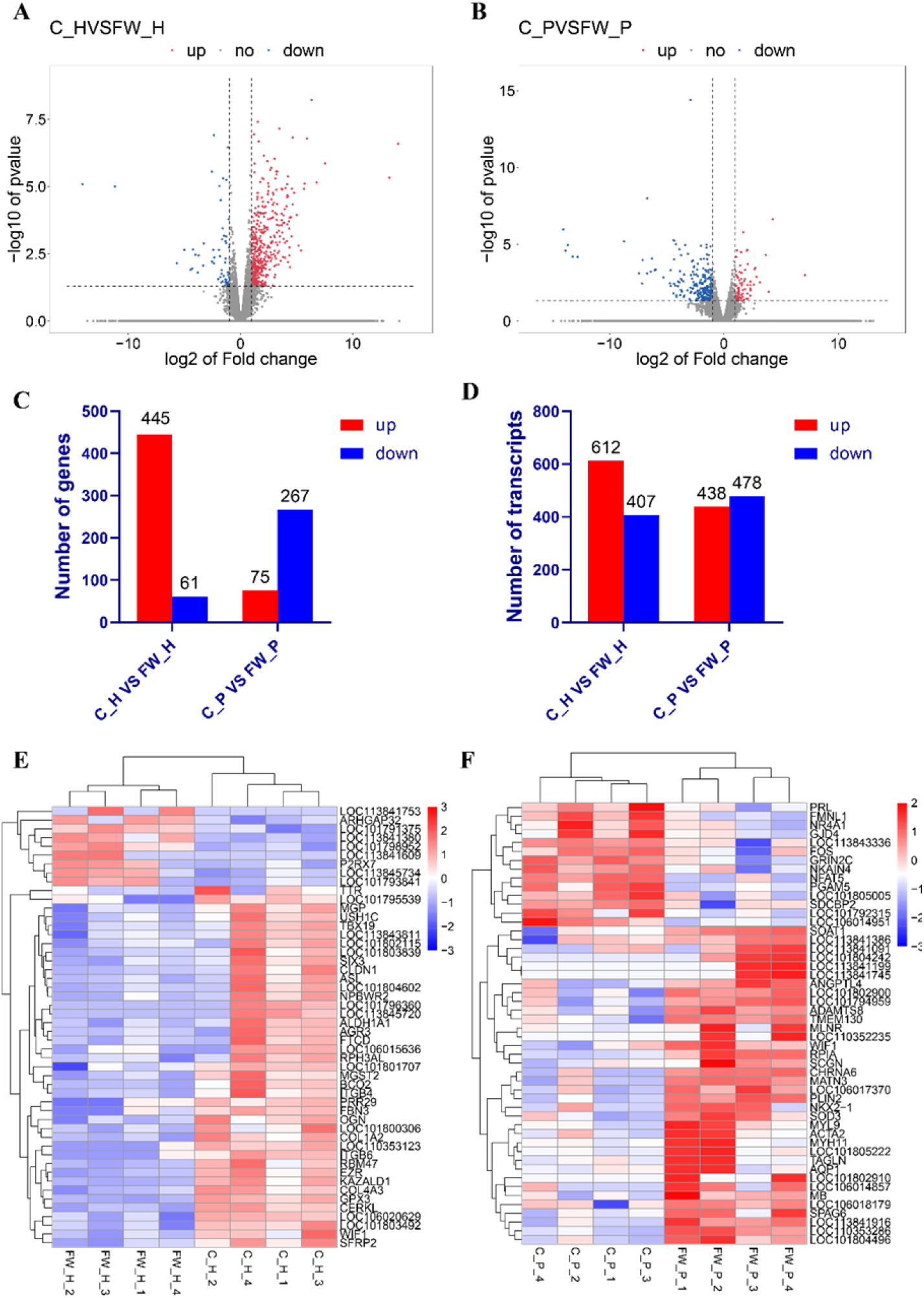
Differential expression analysis of the C and FW groups. (**A, B**) Volcano charts of the genes expressed in the C_H vs. FW_H and C_P vs. FW_P comparisons, respectively. Red dots represent up-regulated DEGs, blue dots depict down-regulated DEGs and gray dots indicate no difference in expression. (**C**) The numbers of up-regulated and down-regulated DEGs in the C_H vs. FW_H and C_P vs. FW_P comparisons. (**D**) The numbers of up-regulated and down-regulated DETs in the C_H vs. FW_H and C_P vs. FW_P comparisons. (**E, F**) Heat maps of DEGs in the C_H vs. FW_H and C_P vs. FW_P comparisons, respectively. The top 50 DEGs are shown in each comparison.

### Enrichment analysis of DEGs

The study then conducted GO enrichment analysis on the DEGs. All the GO terms were classified into three sections, namely; Biological Process (BP), Cellular Component (CC) and Molecular Function (MF), based on functional annotation. The results of GO functional enrichment analysis showed that a total of 378 GO terms were significantly enriched in the C_H vs. FW_H comparison, out of which 259 were enriched in biological process, 50 in cellular component and 68 in molecular function (p < 0.05, Table S5). Notably, the GO terms that might have been associated with acute stress were enriched in the glucocorticoid biosynthetic process, thyroid hormone transport, development of the adrenal gland, positive regulation of cortisol secretion, activity of the corticotropin releasing hormone, the steroid hormone mediated signaling pathway, steroid hormone receptor activity and regulation of response to oxidative stress (Figure 3A, Table S5). Additionally, the significant enrichment of sensory perception of the light stimulus (GO:0050953), suggested that the light in the cage-rearing system elicited a different response from to that in the floor-water combination system. On the other hand, a total of 389 GO terms were significantly enriched in the C_P vs. FW_P comparison, including 290 biological process terms, 34 cellular component terms and 65 molecular function terms (p < 0.05, Table S6).

**Figure 3.**
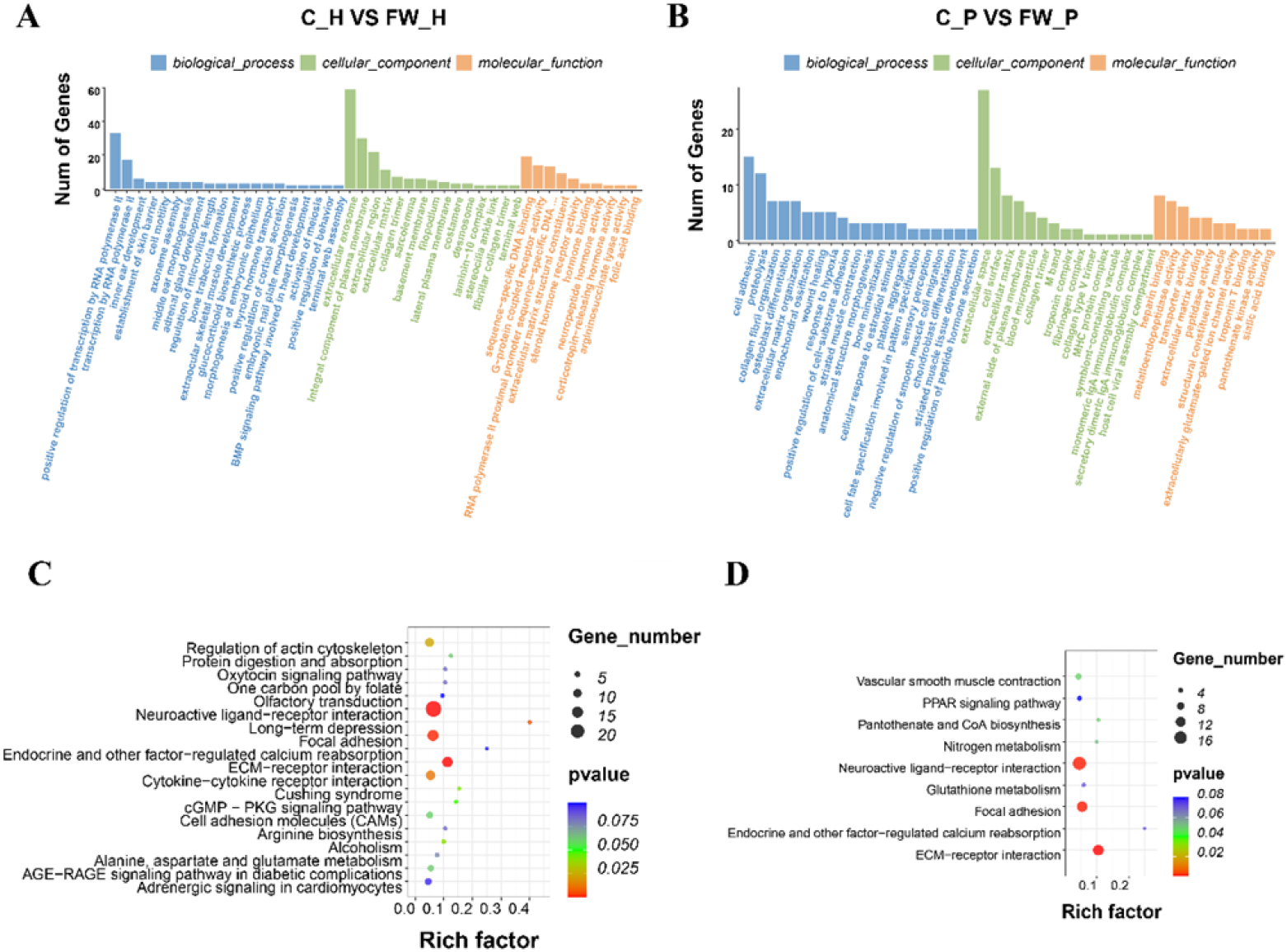
Enrichment analysis of DEGs. (**A, B**) GO classification of the DEGs in C_H vs. FW_H and C_P vs. FW_P, respectively. The top 20, 15, and 10 GO terms in biological processes, cellular components, and molecular functions, respectively, are shown. (**C, D**) The significantly enriched KEGG signaling pathways (P<0.1) of the DEGs in C_H vs. FW_H and C_P vs. FW_P, respectively.

Interestingly, cellular response to the estradiol stimulus, regulation of dopamine secretion, negative regulation of blood pressure, regulation of inflammatory response, detection of oxidative stress and other stress-related terms were among the most significantly enriched GO terms (Figure 3B, Table S6). It is also noteworthy that light associated terms (response to absence of light and retina homeostasis) and reproduction related terms (developmental process involved in reproduction and regulation of female receptivity) were also markedly enriched (Table S6).

Furthermore, KEGG pathway analysis was performed on all the DEGs. Results from the C_H vs. FW_H comparison showed that a total of 19 pathways were significantly enriched (p < 0.1, Figure 3C). The significantly enriched pathways (p < 0.05) are shown in Table 3. Notably, long-term depression, adrenergic signaling in cardiomyocytes and endocrine and other factor-regulated calcium reabsorption were associated with stress (Table S7). Additionally, ECM-receptor interaction, neuroactive ligand-receptor interaction and focal adhesion, were associated with external signal transduction and signaling-molecule interaction. On the other hand, the DEGs in the C_P vs. FW_P comparison were significantly enriched in 9 pathways (p < 0.1, Figure 3D). Interestingly, the top 3 markedly enriched pathways, i.e., ECM-receptor interaction, focal adhesion and neuroactive ligand-receptor interaction, were also among the top KEGG pathways identified in the hypothalamus. Moreover, pathways related to biosynthesis and substance metabolism, including pantothenate and CoA biosynthesis, nitrogen metabolism and glutathione metabolism, were enriched in the C_P vs. FW_P comparison (Table S8). In addition, a total of 13 DEGs were enriched in ECM-receptor interaction pathways, including the Collagen Type VI Alpha 1 Chain (COL6A1), Chondroadherin (CHAD), Thrombospondin 2 (THBS2), Syndecan 1 (SDC1), the Integrin Binding Sialoprotein (IBSP), Tenascin R (TNR), the SLIT and NTRK Like Family Member 1 (SLITRK1), the Von Willebrand Factor (VWF), the Laminin Subunit Alpha 4 (LAMA4), Tenascin C (TNC), the Integrin Subunit Alpha 11 (ITGA11), Glycoprotein Ib Platelet Subunit Alpha (GP1BA) and *COL2A1* (Table 4). Notably, the recurrence of *COL6A2, COL1A2, LAMA5, COL6A1, COL2A1* and *LAMA4* (members of the COL and LAMA families) in both comparisons, suggested that they might be associated with caging related stress in ducks.

**Table 3.** The significantly enriched pathways and enriched genes in the C_H vs.

| Pathway name | p-value | Gene names |
| --- | --- | --- |
| ECM-receptor interaction | 0.000 | *COL6A2, LAMA2, COL1A2, ITGB6, CHAD, COL4A4, COL4A3, SDC1, ITGB4, COL4A5, LAMB1, LAMA5* |
| Neuroactive ligand-receptor interaction | 0.000 | *LEPR, GH1, CGA, PRLHR, PRL, FSHB, TMEM139, NPBWR2, OXTR, MTNR1A, GAL, AVPR1B, GHRHR, TRHR, P2RX7, RXFP1, UCN3* |
| Focal adhesion | 0.001 | *COL6A2, LAMA2, COL1A2, ITGB6, CHAD, COL4A4, COL4A3, ITGB4, MYL9, COL4A5, LAMB1, LAMA5, PDGFRL* |
| Long-term depression | 0.006 | *CRH, LOC113845734* |
| Cytokine-cytokine receptor interaction | 0.012 | *LEPR, GH1, CSF3R, CXCL12, IL17RA, PRL, IL31RA, TNFRSF6B, CXCR6, IL18* |
| Regulation of actin cytoskeleton | 0.024 | *FGF19, IQGAP2, CXCL12, EZR, ITGB6, ITGB4, IQGAP3, MYL9, FGFR4* |
| Alcoholism | 0.040 | *ISLR, CRH, LOC113845734* |
| Cushing syndrome | 0.042 | *CRH, LOC113845734* |
| cGMP - PKG signaling pathway | 0.048 | *MYL9, RGS2* |

**Table 4.** The top 5 enriched KEGG pathways and enriched genes in C_P vs. FW_P

| Pathway name | p-value | Gene names |
| --- | --- | --- |
| ECM-receptor interaction | 0.000 | *COL6A1, CHAD, THBS2, SDC1, IBSP, TNR, SLITRK1, VWF, LAMA4, TNC, ITGA11, GP1BA, COL2A1* |
| Focal adhesion | 0.001 | *COL6A1, CAV3, CHAD, THBS2, IBSP, TNR, MYL9, VWF, PGF, LAMA4, TNC, ITGA11, COL2A1* |
| Neuroactive ligand-receptor interaction | 0.001 | *DRD5, GRM4, GPR50, CHRNA6, GLP1R, GRIA3, SSTR1, NMUR1, KBP, GRIN2C, VIP, CRAMP1, MLNR, MTNR1B* |
| Pantothenate and CoA biosynthesis | 0.049 | *PANK4, PANK3* |
| Vascular smooth muscle contraction | 0.049 | *ACTA2, CALCB, MYL9, KCNMB1* |

### Validation of sequencing data

After conducting enrichment analysis, 5 DEGs were selected from each comparison to validate the sequencing data. Therefore, the study designed the specific qPCR primers for Beta-carotene Oxygenase 2 (BCO2), Matrix Gla Protein (MGP), Loc101791375, Loc106020629 and Corticotropin Releasing Hormone (CRH). The qPCR results showed that the expression trends were consistent with those of the RNA-seq data in the C_H vs. FW_H comparison (Figure 4A). On the other hand, the qPCR-assessed trends in the expression of SOD3, Myosin Light Chain 9 (MYL9), Fos Proto-Oncogene (FOS), Ribose 5-Phosphate Isomerase A (RPIA) and PGAM Family Member 5 (PGAM5) in the C_P vs. FW_P comparison, were consistent with those in the RNA-seq data (Figure 4B).

**Figure 4.**
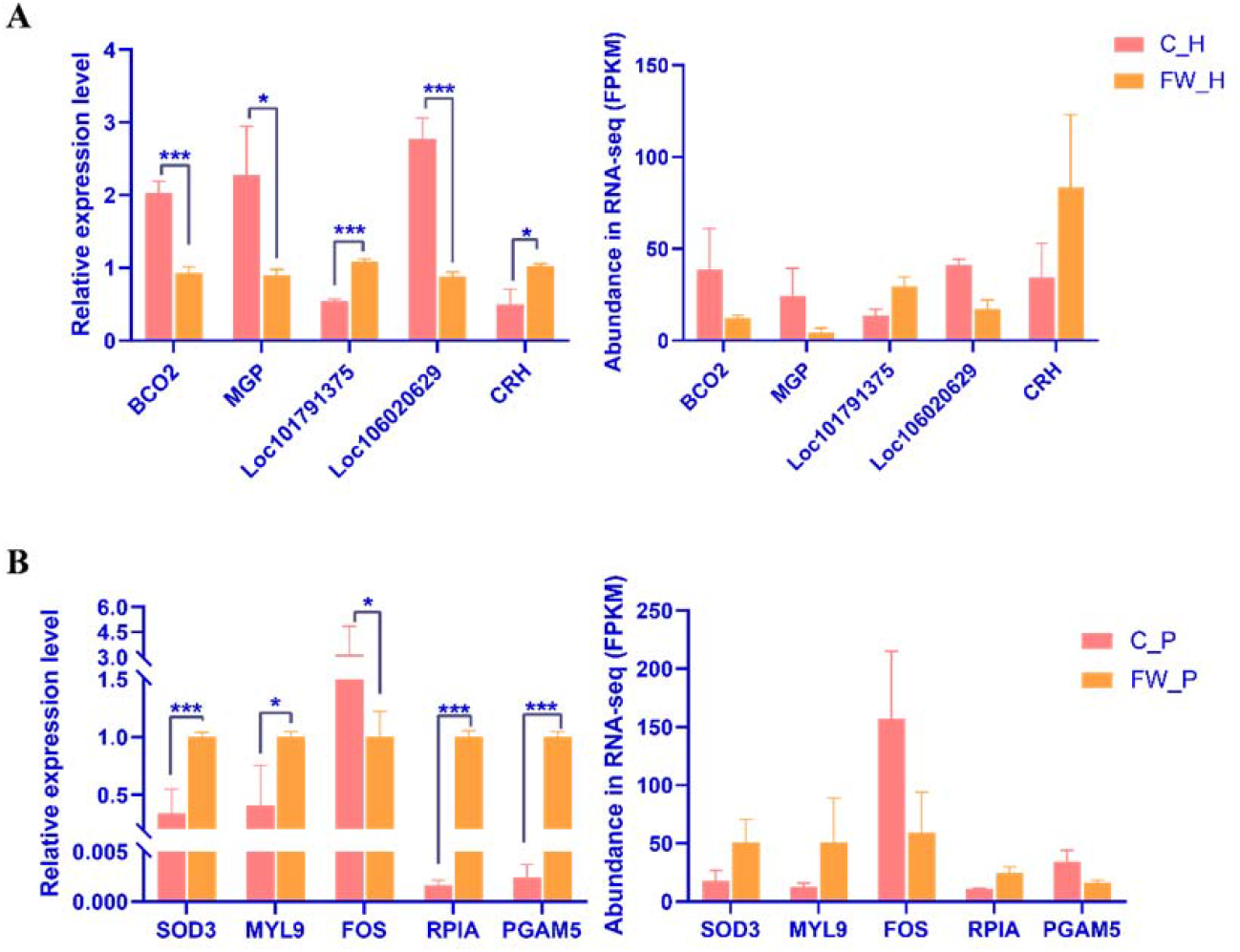
qRT-PCR validation of the DEGs. The graphs on the left represent the qPCR results of the selected DEGs in the (**A**) C_H vs. FW_H comparison and (**B**) in the C_P vs. FW_P comparison. On the other hand, the graphs on the right represent the expression level of DEGs in RNA-seq data. The values are presented as the mean ± SEM in all the panels. * p < 0.05; ** p < 0.01; *** p < 0.001.

### Cage-rearing stress altered RNA splicing

After analyzing the number of differentially expressed transcripts and genes, the results showed that there was a huge difference in the number up-regulated and down-regulated DEGs but little difference between the corresponding up-regulated and down-regulated DETs. Therefore, it was speculated that the parental genes of multiple differentially expressed transcripts were not DEGs. This phenomenon may have occurred because the two differential transcripts from the same parental gene were oppositely regulated or the expression levels of the differential transcripts was lower, which did not significantly affect the expression of the parental gene. In order to verify this hypothesis, the threshold for DETs screening was set at a q-value < 0.05 and the sum of FPKM from 8 samples in one comparison was set to be greater than 20. Based on this threshold, a total of 9 DETs were obtained in the C_H vs. FW_H comparison and 8 DETs met the criteria in the pituitary comparison (Table S9). Notably, there was no difference in the expression of the parental genes of these DETs in their respective comparisons although one or more transcripts were differentially expressed. For instance, in the C_P vs. FW_P comparison, the isoform of Isocitrate Dehydrogenase 1 (IDH1), XM_027461485.1, was significantly down-regulated (q < 0.001) while XM_027461486.1 was up-regulated (p < 0.05). However, there was no difference in the expression of IDH1 between the cage-rearing and the floor-water combination groups (Figure 5A). Similarly, XM_027446641.1 (an isoform of the Receptor Accessory Protein 5, REEP5) was significantly down-regulated (q < 0.05) while XM_027446638.1 was up-regulated (p < 0.05). Nonetheless, there was no difference in the expression of the other two transcripts and the REEP5 gene in the C_P vs. FW_P comparison (Figure 5B). In the hypothalamic comparison, the isoform of the Ribosomal Protein L7a (RPL7A), XM_027470724.1 was significantly down-regulated (q < 0.001) while the expression of the other transcript was significantly higher than that of XM_027470724.1, which did not affect the expression level of the gene (Figure 5C). Moreover, qPCR primers for *IDH1*, *REEP5* and the respective DETs were designed to verify the differential expression of the above transcripts. The results were consistent with the sequencing data, indicating that acute cage-rearing stress can change the alternative splicing of some genes (Figure 5D).

**Figure 5.**
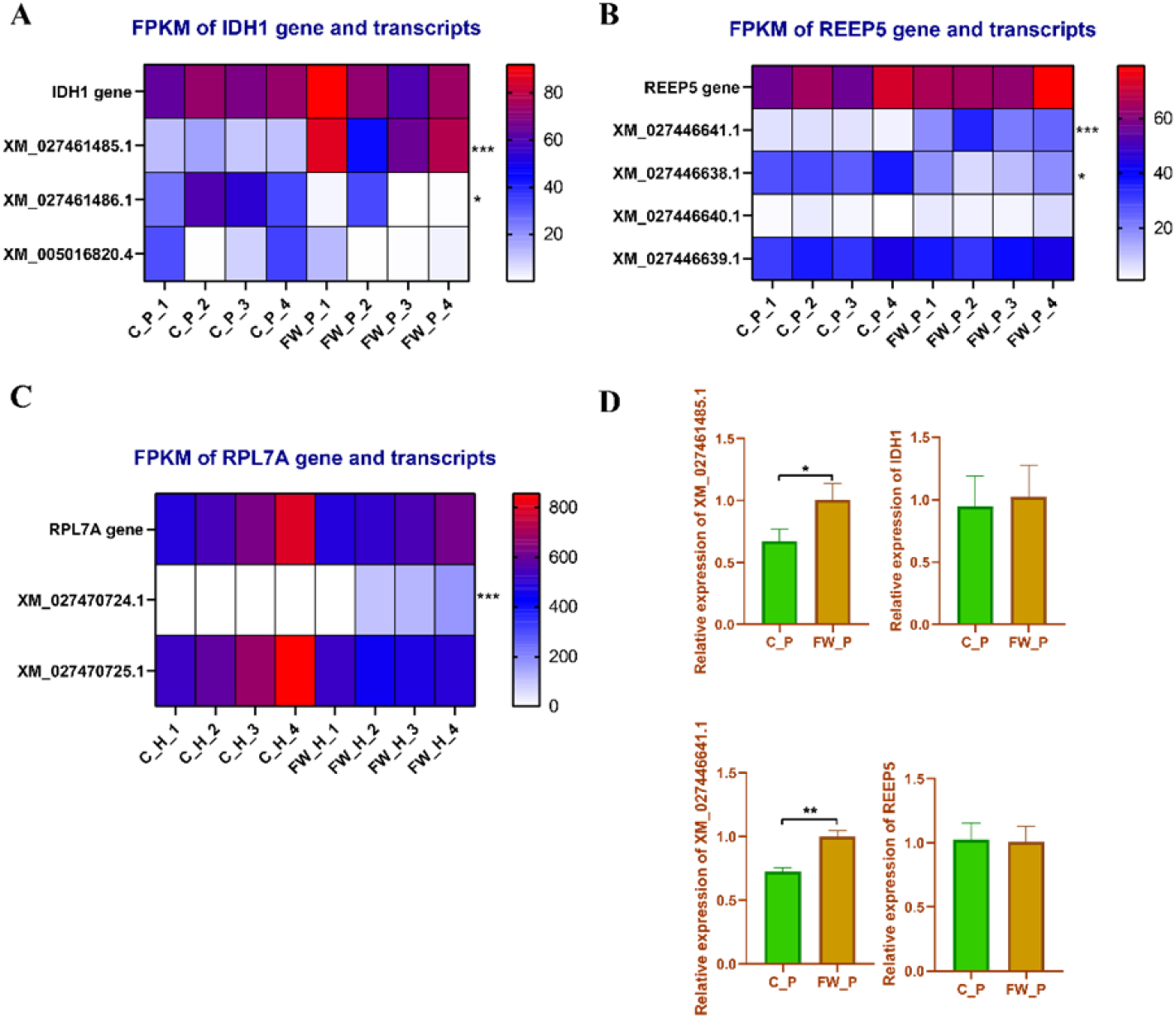
Alternative splicing was different between the C and FW groups. (**A, B**) Heat maps of the expression level of *IDH1, REEP5* and their isoforms in the C_P vs. FW_P comparison. (**C**) A heat map of the expression of *RPL7A* and its transcripts in the C_H vs. FW_H comparison. (**D**) qPCR validation of *IDH1, REEP5* and their isoforms in the C_P vs. FW_P comparison. The values are presented as the mean ± SEM in all the panels. * p < 0.05; ** p < 0.01; *** p < 0.001.

## Discussion

In China, both food safety and environmental protection are given great emphasis, making farmers to explore more environmentally friendly breeding models for egg laying ducks. In addition, methods of duck breeding have gradually changed from the traditional floor-water combination system to cage breeding(Hou and Liu 2021). Compared to a single cage, multilayer cages have a higher feeding density, which allow for better utilization of space and are more suitable for large-scale feeding as well as management. Moreover, multilayer cage-rearing and floor-water combination breeding are two different feeding methods that can be used as models for studying the interaction between organisms and their environment. Consequently, the current study explored the similarities and differences in gene expression between ducks reared in multilayer cages for three days and those raised in the floor-water combination system. The results therefore provide baseline information for future research on the adaptability of ducks to long-term cage-rearing. The study also has an important economic value and practical significance to the waterfowl industry.

Previous research showed that cage-rearing can cause endoplasmic reticulum stress and lead to liver injury in ducks. It was also reported that there was an increase in the expression levels of inflammation-related factors and immune-related genes(Zhang *et al.* 2019). It is noteworthy that changes in the external environment affect the transcriptome of an organism(Krishnan *et al.* 2020; Wang *et al.* 2020; Xu *et al.* 2019). In addition, activation of the HPA axis triggers the hypothalamus to release CRH and Arginine Vasopressin (AVP). CRH stimulates the production and secretion of the Adrenocorticotropic Hormone (ACTH) in the pituitary(Dick and Provencal 2018; Guest and Guest 2018). The present study caged ducks in the floor-water combination system for three days then assessed the differences between the transcriptomes of ducks reared in cages and those raised in the traditional method. According to previous study, cage-rearing stress gradually subsides after the 4^th^ day(Zhang *et al.* 2019). Herein, there was a decrease in the expression of the *CRH* gene although there was no significant difference in the expression of the *ACTH* gene (Proopiomelanocortin, POMC). These indicated that the acute stress in ducks had gradually faded at the three days of caging and that they had adapted to their environment. The up-regulation of *CRH* may have been due to the weak response of cage-reared ducks to capture and sampling. On the other hand, ducks raised in the floor-water combination system had a strong response, leading to the down-regulation of *CRH* and no change in *POMC.*

In addition, the study conducted differential expression analysis between the two groups and enrichment analysis of DEGs after RNA sequencing. The results of enrichment analysis showed that the DEGs were mainly enriched in processes associated with metabolism and processing of environmental information. Notably, the ECM-receptor interaction, neuroactive ligand-receptor interaction and the focal adhesion signaling pathways were the most significantly enriched pathways in both groups. The ECM-receptor interaction signaling pathway belongs to the environmental information processing subclass (https://www.kegg.jp/dbget-bin/www_bget?pathway+hsa04512). The Extracellular Matrix (ECM) is a three-dimensional acellular structure composed of collagen, Proteoglycan (PG) and glycoprotein. The structure mediates cell-matrix or cell-cell adhesion, signal transduction and cell growth(Theocharis *et al.* 2016). In this study, 12 genes including’ *COL6A2, LAMA2, COL1A2* and *ITGB6*, were enriched in the ECM-receptor interaction signaling pathway in the C_H vs. FW_H comparison while 13 genes were enriched in the pituitary. Additionally, collagen is an important member of the ECM-receptor interaction signaling pathway(Vargas *et al.* 2013). The present study showed that several members of the collagen family were significantly differentially expressed. Moreover, it was previously reported that decreasing oxidative stress in cardiomyocytes can reduce the expression of collagen 1 and 3(Zhou *et al.* 2009). In this study, there were significant differences in the expression of *COL6A2, COL1A2, COL4A4*, COL4A3, *COL4A5, COL6A1* and *COL2A1* in the two groups, indicating that they were regulated by stress in the cage environment. Furthermore, the neuroactive ligand-receptor interaction pathway is the aggregation of all receptors and ligands related to intracellular and extracellular signaling pathways in the plasma membrane. The pathway also belongs to the Environmental Information Processing subclass (https://www.kegg.jp/kegg-bin/show_brite?htext=hsa00001.keg&query=hsa04080). Genes in the neuroactive ligand-receptor interaction pathway are associated with stress response, including psychological, electric shock, heat and other forms of stress(Kim *et al.* 2017; Lu *et al.* 2020; Luo *et al.* 2015). In this study, 17 and 14 DEGs were enriched in the neuroactive ligand-receptor interaction pathway in the C_H vs. FW_H and C_P vs. FW_P comparisons, respectively.

It is also worth noting that animals change the levels of certain proteins in the body in response to external environmental stimuli when they enter into different environments(Sapolsky *et al.* 2000). In addition, the body can regulate the expression of some proteins through alternative splicing in order to cope with changes in the external environment(Gopalakrishnan and Kumar 2020; Singh *et al.* 2017; Tapial *et al.* 2017). Alternative splicing is the process of selecting different combinations of splicing sites on an mRNA precursor in order to produce different isoforms, resulting in different phenotypes due to distinct levels of expression in the same cell(Birzele *et al.* 2008). Moreover, previous studies showed that when the body is faced with stress or bacterial infection, it responds by inducing alternative splicing(Martin *et al.* 2019; Staiger and Brown 2013; Sun 2017). In this study, there were no differences in the expression levels of some genes between the cage-rearing and floor-water system groups. However, the expression of certain isoforms changed significantly. These results therefore showed that the caged environment can regulate the body’s response through alternative splicing.

Moreover, light affects the feeding, growth and reproductive behavior of birds(Liu *et al.* 2020; Pitesky *et al.* 2019; Zaguri *et al.* 2020). In the cage-rearing system, poultry receive a stable intensity and duration of light. However, the amount of light received by birds in their natural environment is often unstable due to the influence of weather and geographical location(Rani and Kumar 2014). In this study, the DEGs were enriched in some GO terms related to light perception, including sensory perception of the light stimulus and response to absence of light. This suggested that the different levels of illumination in the cage-rearing and floor-water combination systems affected the growth and development of the ducks.

In this study, the hypothalamus and pituitary of ducks in two different rearing systems were collected for transcriptome sequencing. The results showed that there were significant differences in the expression of stress-related genes and pathways. The findings also showed that the cage-rearing system had an effect on the transcriptome of the ducks. Chronic stress has been the focus of most studies on animal production. Therefore, more attention should be paid to the effects of long-term caging on the growth and production performance of ducks. Additionally, multi-omics can be used to explore the mechanisms underlying the effects of cage-rearing on ducks. Moreover, methods can be established to not only mitigate the impact of caging but also enhance the growth and reproductive performance of ducks.

## Conclusions

In summary, the present study assessed the differences in the expression of genes between ducks in the cage-rearing and floor-water combination systems. The results showed that there was a significant change in the expression of genes after short-term caging. The findings also revealed that pathways associated with endocrine and environmental information processing were significantly enriched. Notably, ECM-receptor interaction, neuroactive ligand-receptor interaction and focal adhesion were the main signaling pathways enriched in response to short-term caging. Additionally, the caged environment can regulate the body’s response through alternative splicing. These results can therefore help in understanding the mechanisms underlying the effect of cage-rearing on the growth and reproduction of waterfowl. The findings also highlight the gene regulatory networks involved in animal responses to acute stress.

## Acknowledgments

This study was support by the Technology System of Modern Agricultural Poultry Industry of Jiangxi Province (JXARS-09), Research and optimization of shell-less hatching method of avian embryos (190223) and Doctoral start-up fund of Jiangxi Agricultural University. The first author thanks Mrs. Jing Liu and Zhuo Chen for their assistance.

## Data Availability Statement

All sequencing data were uploaded to in the Gene Expression Omnibus (GEO) with accession number GSE173134 (https://www.ncbi.nlm.nih.gov/geo/query/acc.cgi?acc=GSE173134).

